# An improved *Trypanosoma vivax* genome assembly confirms the distinct organisation of its variant antigen repertoire

**DOI:** 10.64898/2026.10.08.757617

**Authors:** Andrew P. Jackson, Rachel Young, Mark A. Whitehead, Pieter C. Steketee, Alistair C. Darby, Liam J. Morrison

## Abstract

*Trypanosoma vivax* is a unicellular parasite spread by biting flies and an important livestock pathogen across sub–Saharan Africa and South America. Alongside its better known relative, *T. brucei*, it causes animal trypanosomosis. An accurate genome sequence is essential to functional genetics as well as the design of novel diagnostics, drugs and vaccines. However, the existing *T. vivax* genome sequence is fragmentary, making it difficult to examine genomic features such as the Variant Surface Glycoprotein (VSG) genes that are the basis for antigenic variation, a key phenotype that underpins immune evasion by African trypanosomes. Our aim was to produce a *T. vivax* genome sequence with improved contiguity to properly characterise the VSG genes. We present a new core genome assembly in 23 scaffolds, forming approximately 10 chromosomes. Many scaffolds are articulated with extensive sub-telomeric domains that, like *T. brucei*, contain VSG gene arrays but in *T. vivax* these are near-perfect repeats of intact genes. Using transcriptomic and proteomic analysis of VSG expression during bloodstream infection we show that the active VSG is expressed from a sub-telomeric array while other arrays are silent. We find no evidence for a VSG expression site at the telomeres or elsewhere. Therefore, although monoallelic expression and antigenic switching occur in *T. vivax*, the familiar regulatory mechanism of *T. brucei* does not exist in *T. vivax*. This improved *T. vivax* genome assembly demonstrates that *T. brucei* is a poor guide for *T. vivax* genetics generally and for regulation of antigenic variation specifically.

**Significance statement:** *Trypanosoma vivax* is a parasite spread by biting flies and the cause of an important livestock disease, African animal trypanosomosis, across sub–Saharan Africa and South America. Like all African trypanosomes, including the model organism *T. brucei, T. vivax* evades host immunity through antigenic variation of its Variant Surface Glycoprotein (VSG) genes. The existing *T. vivax* genome sequence is too fragmentary to understand the regulation of VSG expression. Here, we present a new genome assembly of approximately 10 chromosomes. We show that, unlike *T. brucei, T. vivax* VSG are arranged in tandem arrays of intact VSG genes, the active VSG is expressed from one tandem array while other arrays are silent, and there is no evidence for a telomeric VSG expression site. Therefore, although monoallelic expression and antigenic switching occur in *T. vivax*, the regulatory paradigm familiar from *T. brucei* does not apply and must be different.

## Introduction

African Animal trypanosomosis (AAT) is a vector-borne disease of livestock caused by multiple species of African trypanosome (*Trypanosoma* spp.). Among these, the zoonotic pathogen *T. brucei* is well studied, not least for the sophisticated antigenic variation of its Variant Surface Glycoproteins (VSG) (Mugnier et al. 2016; Sima et al. 2019). *T. vivax* is less familiar but is among the most frequent and important causes of AAT (Fetene et al. 2021; Osório et al. 2008). Across sub-Saharan Africa, AAT is transmitted by tsetse flies (*Glossina* spp.) but has also spread to parts of tropical Asia and South America where it is spread mechanically by other biting flies (Cecchi et al. 2015). In Africa, AAT is endemic in 33 countries where it has a profoundly negative effect on livestock production through increased animal morbidity and mortality, as well as reduced land use, and therein on socio-economic development (Swallow 2000; Tora & Dana 2024; Adane et al. 2025). The cost of AAT is approximate but measured in billions of dollars per year (Kristjansen et al. 1999; Abro et al. 2021), and disease elimination is typically predicted to double agricultural productivity across Africa (Shaw 2009; Shaw et al. 2014, 2015; Okello et al. 2015). Presently, improvements in pathogen biology required to develop novel interventions for AAT are limited by the lack of molecular resources and tools (Morrison et al. 2016).

An accurate genome sequence is the starting point for parasite genetics, molecular biology, epidemiology and immunology. Compared to the highly contiguous, chromosome-level *T. brucei* genome assembly, the *T. vivax* Y486 genome sequence is underdeveloped. The current reference sequence was produced using Sanger technology and scaffolded on the 11 *T. brucei* chromosomes (Jackson et al. 2012); the true *T. vivax* karyotype remains uncertain. The current sequence has shown that *T. vivax* has various species-specific genes, especially those predicted to expressed on the cell surface (Greif et al. 2013; Jackson et al. 2013). Transcriptomic and proteomic studies have confirmed that *T. vivax* has distinct developmentally regulated cell surface proteins to *T. brucei* (Greif et al. 2013; Jackson et al. 2015), including one (Romero-Ramirez et al. 2022) that has potential as a vaccine candidate (Autheman et al. 2021).

All African trypanosomes display antigenic variation (AV) of a homologous Variant Surface Glycoprotein (VSG), which is crucial to their host-parasite interactions, immunity and epidemiology. The seminal comparison of African trypanosome genomes showed how the VSG genes underpinning AV have distinct evolutionary histories, which indicated that their regulation was different despite the common phenotype (Jackson et al. 2012). In *T. brucei*, VSG are arranged in irregular arrays of pseudogenes and gene fragments within sub-telomeres of megabase chromosomes as well as small, accessory mini-chromosomes (Berriman et al. 2005). VSG transcription is restricted to telomeric expression sites and VSG are activated after switching into an expression site through gene conversion, which also leads to assortment of VSG sequences, and an almost unlimited supply of antigenic diversity (Faria et al. 2022). In a previous population genomic study, we showed that *T. vivax* VSG sequences are not assorted by gene conversion and antigenic diversity is quite limited by comparison (Silva Pereira et al. 2020). Moreover, it is known that *T. vivax* lacks mini-chromosomes (Van der Ploeg et al. 1984) and its VSG loci appear to be mostly intact (Jackson et al. 2012), but the current assembly does not allow a comprehensive analysis. Further progress on understanding AV in *T. vivax* therefore requires an accurate genome sequence.

In this study, we present a revised *T. vivax* Y486 genome sequence and annotation with telomere-to-telomere assembly of several chromosomes. Our aims are to 1) compare the chromosomal order of *T. vivax* with *T. brucei* to measure the extent of conserved homoeology; 2) catalogue the number and positions of VSG loci; 3) search for a VSG gene cassette at telomeres or elsewhere that is analogous to the *T. brucei* expression site; and 4) measure VSG gene and protein expression in Y486 to identify the genomic positions of expressed VSG.

## Results and Discussion

### An improved *T. vivax* reference genome assembly

The existing *T. vivax* Y486 genome sequence comprises 47.5 Mbp and 8,290 contigs (N50 = 21.8 kbp) formed into pseudochromosomes based on homology with the 11 *T. brucei* 927 megabase chromosomes (Jackson et al. 2012). We produced a new assembly from a combination of Hifi (PacBio) and Novaseq (Illumina) data, scaffolded with HiC reads. The initial contig assembly based on hifi reads only was 84.6 Mbp in length (124 contigs, N50 = 2.85 Mbp). Following two rounds of haplotig purging the genome was assembled using Hi-C reads with YaHS (v1.2a.2). Assembly size was reduced to 70.7 Mbp (29 contigs, N50 = 3.65 Mbp). This sequence is 100% complete based on BUSCO analysis of 130 genes in the Euglenozoa_odb10 database (Manni et al. 2021).

DNA contacts imputed from Hi-C read mapping are shown in Fig. S1. This shows that there are 18 genomic domains with enriched contacts that are suggestive of chromosome-sized scaffolds (‘sc’). However, there are also abundant contact points between these putative chromosomes, which we interpret as interactions between the sub-telomeric domains known to exist in this genome, which are enriched with transposable elements, paralogous gene families and other repetitive material (Jackson et al. 2012).

Adding Illumina reads to the HiFi-Hi-C assembly did not deliver any improvement; however, we also assembled these short reads alone to check that small DNA species had not been omitted from the assembly by size-selection while preparing the DNA sequence library. This produced a highly fragmentary assembly, (78,442 contigs, N50=4,336 bp), as expected. After excluding >14,000 mtDNA minicircles, there were 53 contigs >1 Kbp in length, which could not be mapped to the HiFi-Hi-C assembly. Some of these contigs contain additional VSG genes, (though not other sequences), and so they were appended to the HiFi-Hi-C assembly as a contig ‘bin’.

Comparing the 29 contigs produced by the HiFi-Hi-C assembly to each other using BLASTn identified several regions of near-perfect identity between contigs ranging between 4 and 70 Kbp in length. After accepting these manual articulations, the assembly includes 7 complete chromosomes (sc1+9+19, 8+13+17, 2, 5+15, 6+20, 3 and (tentatively) 7+14). Then another seven incomplete chromosomes with only one telomere (sc4+10, 11, 12, 16, 18, 21 and 22). There are another five small contigs with no genes on them (sc24, 26, 27, 28 and 29), two of which (24 and 29) have telomeres. Two original contigs (23 and 25) were identical to larger contigs so are purged as alternative haplotypes.

Resolving these sequence overlaps reduced the final assembly to 24 scaffolds; all but the smallest of these are shown in Fig. 1. These include seven complete chromosomes (sc1-6 and sc8) with a telomere-to-telomere assembly, four of which have alternative sub-telomeres (sc1a, 2a, 3a and 5a). This appears to emulate *T. brucei* chromosomes, which typically have up to four hemizygous sub-telomeric domains for each distinct chromosomal core (Müller et al. 2018). Another six chromosome-sized scaffolds have only one telomere (sc7, 9-13); these contain non-overlapping, ‘core’ polycistrons, and so are unlikely to be alternative sub-telomeres of other chromosomes and we expect them to ultimately resolve into three chromosomes. Scaffold 7a, by contrast, contains largely sub-telomeric material and overlaps with sc7. A further five small scaffolds contain either isolated telomeres (sc14 and sc15) or no genes or telomeres (sc16-18). Scaffold 19 is the mtDNA maxicircle. Overall, we propose that *T. vivax* is likely to have 10 megabase chromosomes.

**Fig. 1.**
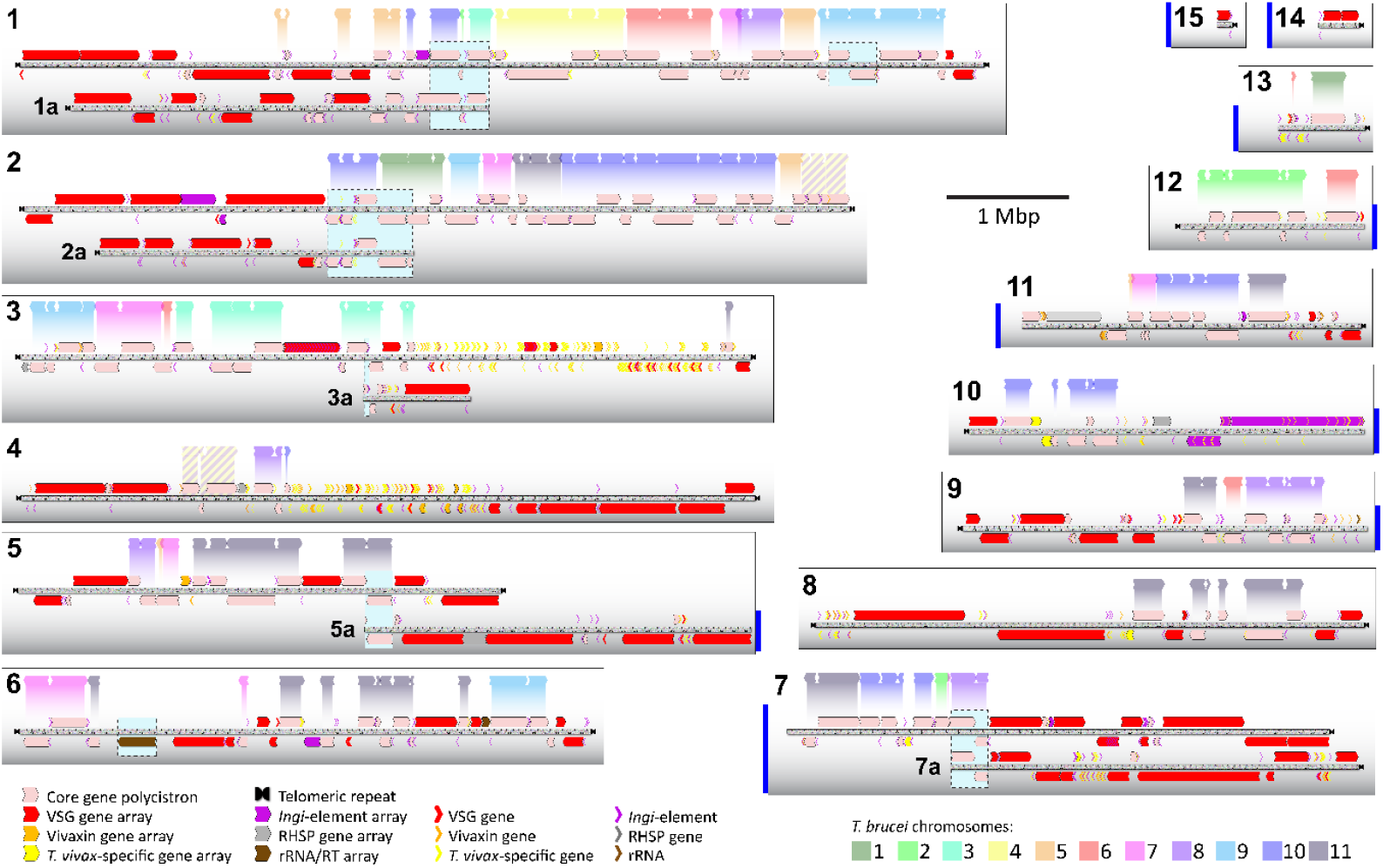
A *Trypanosoma vivax* Y486 hybrid HiFi-Hi-C genome assembly. Chromosomal scaffolds (sc1-15) are represented by scaled grey bars. Polycistronic transcription units (pink) and other features are shown by coloured arrows. Sc1-6 & 8 are complete chromosomes, often with proposed alternative sub-telomeres (marked ‘a’). Sc7,9-15 are partial chromosomes; vertical blue bars indicate unclosed gaps. Conserved synteny with *T. brucei* chromosomes (colour-coded) is indicated above each scaffold.

We found no evidence for accessory chromosomes in the assembly, corroborating the original position of Van der Ploeg et al. (1984). The mini-chromosomal sequence proposed by Dickin & Gibson (1989) based on a repeat satellite can be located on scaffold 10 (1.68-1.78 Mbp).

We attempted to close the persistent assembly gaps to produce complete chromosome sequences. We generated ONT read data from the same gDNA preparation and trialled multiple assembly and hybrid-assembly approaches to incorporate the different datasets (ONT only, Hi-C + ONT, HiFi + Hi-C + ONT) and tools (Kolmogorov et al. 2019), Verkko (Antipov et al. 2025), hifiasm and MaSuRCA (Zimin et al. 2013)). ONT-only assemblies for hifiasm and Flye failed to complete. Verkko and MaSuRCA yielded fragmented assemblies after being scaffolded with Hi-C. An assembly combining HiFi and ONT data using hifiasm completed and was of comparable contiguity to the HiFi-Hi-C assembly (i.e. 68.16 Mbp; N50 = 4.68 Mbp). However, the Hi-C contact map reveals that some contigs lack internal Hi-C contacts, suggesting the incorporation of multiple chromosomes into single scaffolds (Fig. S2). The longest assembled contig was 11.2 Mbp, larger than the longest contigs in *T. brucei* (chr11, 5.6 Mbp; Müller et al. 2018) or *T. congolense* assembles (‘tco13a’, 8 Mbp; Krasiļņikova et al. 2026). Furthermore, this and other large scaffolds contained internal telomeric repeats. We therefore decided the ONT data failed to improve assembly and utilised only the HiFi-Hi-C assembly.

### *T. vivax* and *T. brucei* both have ‘fused’ chromosomes but share no homoeology

The general organisation of the *T. vivax* genome is superficially similar to *T. brucei*. Housekeeping genes are contained within polycistrons that collectively form chromosomal ‘cores’; these are shown in pink in Fig. 1 and are colinear with their homoeologs in *T. brucei*; indeed, they are conserved across the Trypanosomatidae (El-Sayed et al. 2005). Between these cores and the telomeres are large, sub-telomeric regions that contain VSG and other species-specific genes. As in *T. brucei*, these sub-telomeres appear to be hemizygous, i.e. two versions of each sub-telomere are present, but these have no homology. The chromosomes are larger but fewer than is typical for other trypanosomatids (Almutairi et al. 2021; Davey et al. 2021; Greif et al. 2026), showing that, like *T. brucei, T. vivax* chromosomes originated through the fusion of ancestral chromosomes.

Until now, it was not possible to examine karyotypic evolution because the *T. vivax* sequence was scaffolded against *T. brucei*. A recent chromosomal comparison of *T. brucei* with its closer relative *T. congolense* showed numerous rearrangements (Krasiļņikova et al. 2026). Unsurprisingly, alignment of *T. vivax* and *T. brucei* shows that there is no substantial homoeology between their karyotypes; each *T. vivax* chromosome contains regions of homology to multiple *T. brucei* chromosomes (Fig. 1). As with previous trypanosomatid genome comparisons, the integrity of polycistrons is maintained and breakpoints in co-linearity coincide with strand switch regions or transposon insertion points (El-Sayed et al. 2005). Thus, while there is local co-linearity of gene order within polycistrons, there is no homoeology of entire chromosomes. Hence, it is not possible to say whether *T. vivax* and *T. brucei* each inherited already fused chromosomes that have subsequently been reorganized or independently fused ancestral chromosomes that were smaller and more numerous. What is clear is that the *T. brucei* assembly offers no guide for scaffolding *T. vivax* chromosomes.

As with *T. brucei*, the *T. vivax* genome possesses a 0.5 Mbp segmental duplication, (striped yellow and purple on scaffolds 2 and 4 in Fig. 1), which corresponds with chromosome 31 in *Leishmania* spp., recently characterised as the trypanosomatid ancestral supernumerary chromosome (TASC; Reis-Cunha et al. 2024). The two duplicons have diverged through extensive gene loss (and some additions) since duplication as was observed in *T. brucei* (Jackson 2007).

Like other African trypanosome genomes, *T. vivax* has large sub-telomeres containing contingency genes, but there the familiarity ends. Besides its VSG (see below), *T. vivax* possesses many putative cell-surface proteins that are absent from both *T. brucei* and *T. congolense*. We previously arranged these into multigene families (Fam27-44; (Jackson et al. 2013)) and showed that most are developmentally regulated (Jackson et al. 2015). The largest family (Fam34) encodes highly abundant (but not antigenically variant) bloodstream-stage surface proteins expressed alongside the VSG, which we called ‘vivaxin’ (Romero-Ramirez et al. 2022). Fig. 1 shows that vivaxin and other *T. vivax*-specific surface gene families are concentrated in two sub-telomeric regions (on sc3 and 4), where they are arranged in irregular arrays. These arrays seem to have expanded through segmental duplications and 220/489 (45%) of these coding sequences are predicted to be pseudogenes. It may be that much of the apparent diversity of these families may not be expressed, as appears to be the case for vivaxin (Romero-Ramirez et al. 2022)).

### VSG genes are organised into tandem arrays of near-perfect repeats

VSG genes were located using tBLASTn and query sequences representative of the 178 VSG phylotypes previously identified from diverse *T. vivax* strains (Silva-Pereira et al. 2020). In total, 2,666 VSG genes were identified at 151 loci (see Table S1). Most VSG genes occur in sub-telomeric tandem arrays (2,610/2,666 genes; 85/141 loci), which have a typical arrangement. Each array is a near-perfect repeat of the VSG coding region and intergenic sequence and is often flanked by repetitive regions containing ingi transposable elements at the 5’ (55/85 loci) and 3’ ends (51/85 loci), unless adjacent to a telomere. Although there is polymorphism within an array, with VSG tandem copies displaying 1-2% variation at the amino acid level, we consider each array to encode a single antigen. Each array encodes a single VSG phylotype, as defined previously (Silva-Pereira et al. 2020), and each phylotype is only found in one array. There are 56 VSG genes not found in arrays, belonging to 12 phylotypes (e.g. P2, P40, P142; see Table S1). These singleton VSG are located throughout the genome, often adjacent to unrelated *T. vivax*-specific gene families and have multiplied through the same segmental duplications mentioned above.

Having realised that most VSG genes are arrayed in near-perfect repeats, it became clear that there were genes in 26 sub-telomeric arrays (whose locations are noted in Table S1) that represented additional phylotypes but had not previously been annotated as VSG. These were omitted before because they lack primary structural homology to known VSG and so are not recovered by BLAST. We confirmed that these genes encode proteins consistent with the size (350-450 amino acids) and expected tertiary structure (i.e. three-helical bundle) of all known VSG.

### No evidence for an analogous VSG expression site

We have shown previously that most of the *T. brucei* expression site-associated genes are genetic lineages that evolved in *T. brucei* only (Jackson et al. 2013). Hence, we should not expect a VSG expression site containing coding sequences homologous to ESAGs to exist at *T. vivax* telomeres. In *T. congolense*, a gene cassette containing a VSG, hyper-conserved non-coding sequences and non-VSG genes possibly analogous to ESAGs, is repeated at the telomeres of accessory chromosomes (Abbas et al. 2018; Krasiļņikova et al. 2026). *T. vivax* might possess a similar, but strictly analogous, site for VSG expression. We examined all peri-telomeric sequences for a repeated gene cassette or hyper-conserved non-coding sequence that could represent such a site but found none. Indeed, although around 19 VSG arrays are found adjacent to telomeres in *T. vivax*, they are only orientated correctly for expression in six cases (see Table S1). We found no regular sequence pattern to suggest that these are expression sites, or different to other arrays in any way.

We then examined the entire genome for repeated gene cassettes, in case the expression site was not telomeric. This did identify dispersed segmental duplications of coding sequences, but none were associated exclusively with VSG arrays. The most abundant example of a gene cassette repeated around the genome concerns a 210 Kbp segmental duplication that is found in 16 locations across 12 scaffolds and includes 11 genes. The variable size and completeness of the region, which sometimes is found flanking VSG arrays is described in Fig. S3. We consider this dispersed, segmental duplication to be the result of the coding region being copied and transposed during sub-telomeric rearrangements, rather than representing *T. vivax* expression site-associated genes. Since VSG arrays are common features of sub-telomeres, this would account for the apparent association.

### Active VSG genes are expressed from sub-telomeric arrays

As the genome sequence offers no evidence for a dedicated VSG expression site, we sought to establish what the active VSG(s) in this Y486 strain are and where in the genome they are encoded. We infected mice with the same Y486 stabilate that was used to prepare the genome sequence and generated a transcriptome and proteome from parasites at the first peak of parasitaemia. RNAseq reads mapped to 1,341 VSG genes (see Table S2) but most of these VSG transcripts (1,281/1,341) had negligible abundance (TPM<1). Fig. 2a shows that there were two super-abundant VSG transcripts TvY486_LIV26_0061645 (TPM=2699.6) and TvY486_LIV26_0013985 (TPM=1625.3), which belong to P151 (scaffold 7a; array 88) and P36 (scaffold 1a; array 11), respectively. These two VSG were among the most numerous transcripts in the cell, comparable to those encoding histones and ribosome components. Fig. 2b shows that these VSG transcripts comprised 12.8% of all transcripts when using ONT reads (although only 1.2% when using Novaseq reads, possible due to short reads being removed due to multi-mapping). Moreover, Fig. 2c shows that they were also an order of magnitude more abundant than the next most abundant VSG (TvY486_LIV26_0017450; TPM=436.2). For these reasons, we consider P151 and P36 to be the major clones, or active VSG during first peak of the infection.

**Fig. 2.**
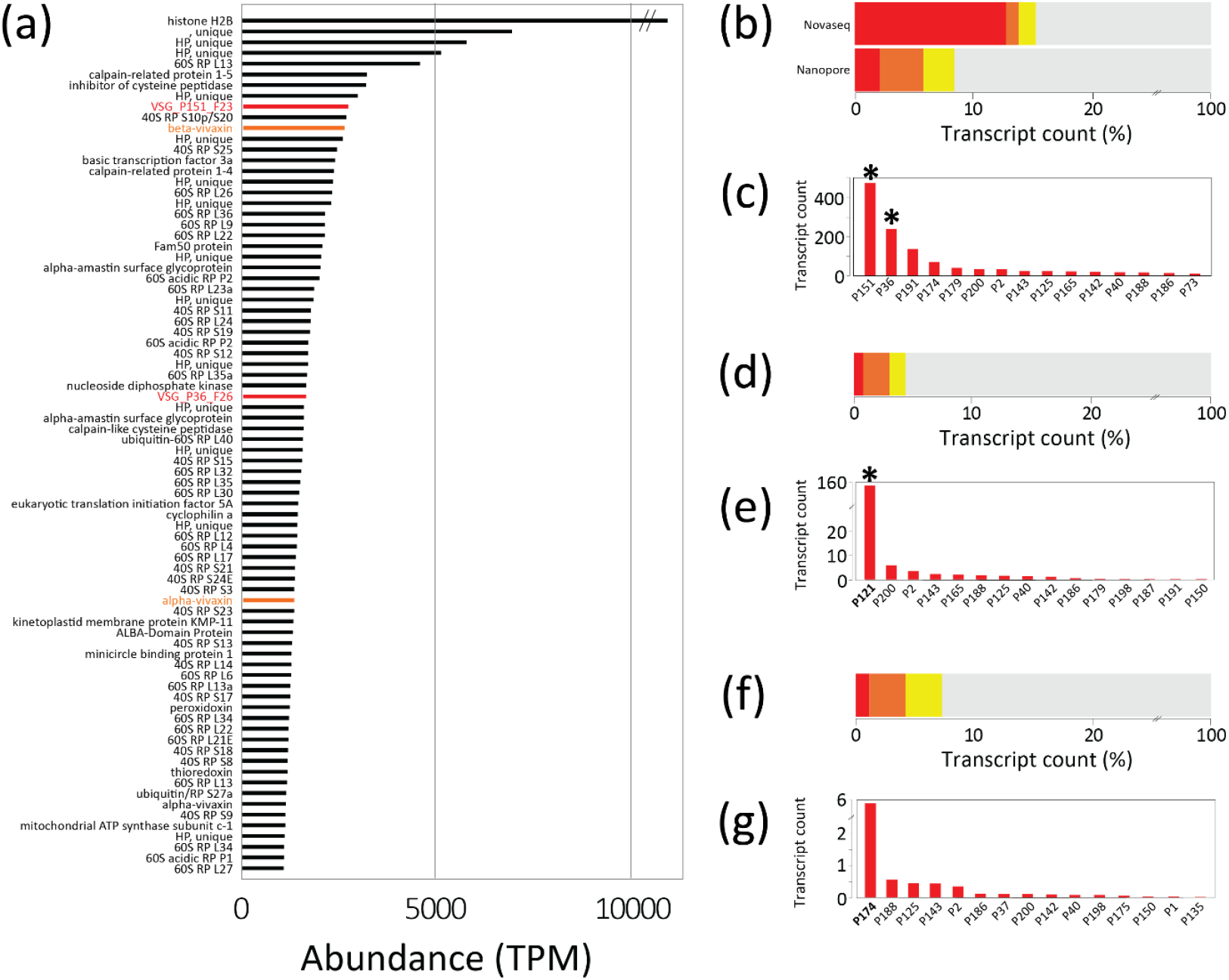
Transcriptomic analysis of VSG expression during a *Trypanosoma vivax* Y486 bloodstream infection in a mouse model compared. **(a)** A bar chart showing the most abundant transcripts, (top 80 only), as determined by mapping RNA-seq reads to all coding sequences with Kallisto; VSG (red) and vivaxin (orange) are highlighted. HP – hypothetical protein. RP – ribosomal protein. **(b)** A stacked bar chart showing the percentage of transcripts deriving from VSG (red), vivaxin (orange) and other *T. vivax* surface gene families (yellow) relative to all other genes (grey). The same RNA sample was sequenced on both Novaseq and ONT platforms for comparison. **(c)** A histogram showing the summed transcript abundance of VSG phylotypes, (top 15 only), as determined by FeatureCounts. **(d-e)** A comparison with data from a previous mouse infection with *T. vivax* Y486 (Jackson et al. 2015). **(f-g)** A comparison with data from a previous goat infection with *T. vivax* Lins (Silva-Pereira et al. 2020). An asterisk ^*^ denotes that the VSG transcript was observed in proteomic analysis (see Table 1, Table S3).

**Table 1.**
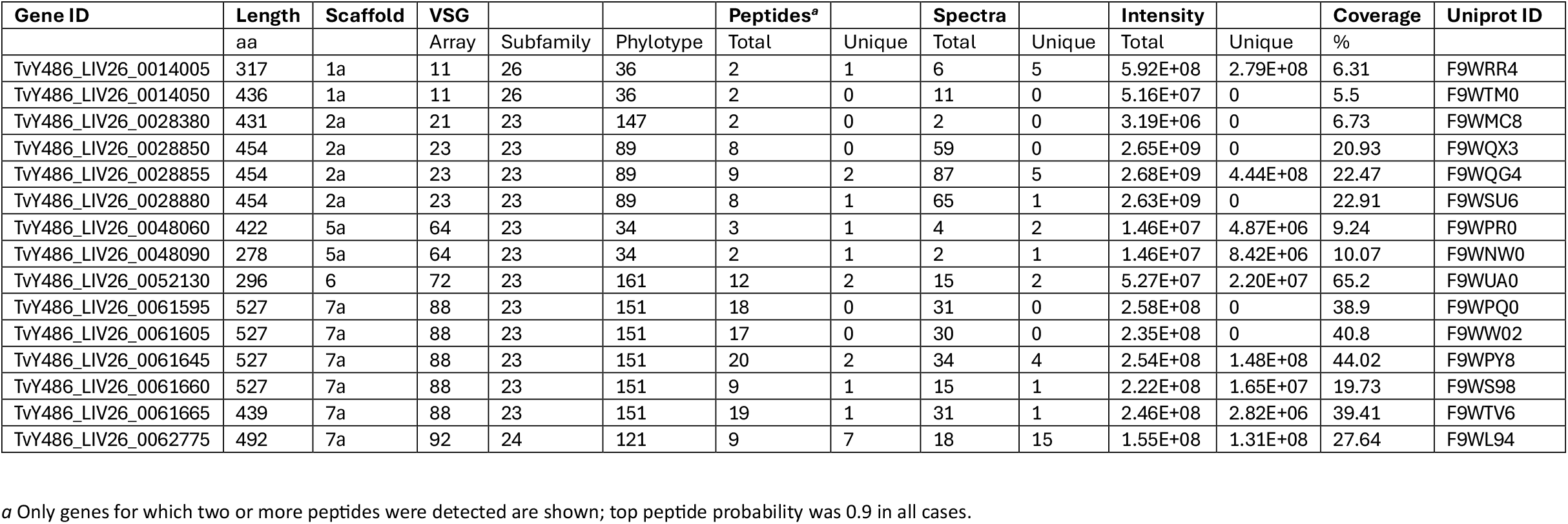
Expressed VSG genes detected in proteomic analysis of a *T. vivax* Y486 blood infection in a mouse model.

| Gene ID | Length | Scaffold | VSG |  |  | Peptides<sup>a</sup> |  | Spectra |  | Intensity |  | Coverage | Uniprot ID |
| --- | --- | --- | --- | --- | --- | --- | --- | --- | --- | --- | --- | --- | --- |
|  | aa |  | Array | Subfamily | Phylotype | Total | Unique | Total | Unique | Total | Unique | % |  |
| TvY486_LIV26_0014005 | 317 | 1a | 11 | 26 | 36 | 2 | 1 | 6 | 5 | 5.92E+08 | 2.79E+08 | 6.31 | F9WRR4 |
| TvY486_LIV26_0014050 | 436 | 1a | 11 | 26 | 36 | 2 | 0 | 11 | 0 | 5.16E+07 | 0 | 5.5 | F9WTM0 |
| TvY486_LIV26_0028380 | 431 | 2a | 21 | 23 | 147 | 2 | 0 | 2 | 0 | 3.19E+06 | 0 | 6.73 | F9WMC8 |
| TvY486_LIV26_0028850 | 454 | 2a | 23 | 23 | 89 | 8 | 0 | 59 | 0 | 2.65E+09 | 0 | 20.93 | F9WQX3 |
| TvY486_LIV26_0028855 | 454 | 2a | 23 | 23 | 89 | 9 | 2 | 87 | 5 | 2.68E+09 | 4.44E+08 | 22.47 | F9WQG4 |
| TvY486_LIV26_0028880 | 454 | 2a | 23 | 23 | 89 | 8 | 1 | 65 | 1 | 2.63E+09 | 0 | 22.91 | F9WSU6 |
| TvY486_LIV26_0048060 | 422 | 5a | 64 | 23 | 34 | 3 | 1 | 4 | 2 | 1.46E+07 | 4.87E+06 | 9.24 | F9WPR0 |
| TvY486_LIV26_0048090 | 278 | 5a | 64 | 23 | 34 | 2 | 1 | 2 | 1 | 1.46E+07 | 8.42E+06 | 10.07 | F9WNW0 |
| TvY486_LIV26_0052130 | 296 | 6 | 72 | 23 | 161 | 12 | 2 | 15 | 2 | 5.27E+07 | 2.20E+07 | 65.2 | F9WUA0 |
| TvY486_LIV26_0061595 | 527 | 7a | 88 | 23 | 151 | 18 | 0 | 31 | 0 | 2.58E+08 | 0 | 38.9 | F9WPQ0 |
| TvY486_LIV26_0061605 | 527 | 7a | 88 | 23 | 151 | 17 | 0 | 30 | 0 | 2.35E+08 | 0 | 40.8 | F9WW02 |
| TvY486_LIV26_0061645 | 527 | 7a | 88 | 23 | 151 | 20 | 2 | 34 | 4 | 2.54E+08 | 1.48E+08 | 44.02 | F9WPY8 |
| TvY486_LIV26_0061660 | 527 | 7a | 88 | 23 | 151 | 9 | 1 | 15 | 1 | 2.22E+08 | 1.65E+07 | 19.73 | F9WS98 |
| TvY486_LIV26_0061665 | 439 | 7a | 88 | 23 | 151 | 19 | 1 | 31 | 1 | 2.46E+08 | 2.82E+06 | 39.41 | F9WTV6 |
| TvY486_LIV26_0062775 | 492 | 7a | 92 | 24 | 121 | 9 | 7 | 18 | 15 | 1.55E+08 | 1.31E+08 | 27.64 | F9WL94 |

Since VSG arrays contain highly similar tandem gene copies, all of which belong to the same phylotype, multiple genes may contribute to the active VSG. Previously, in an experimental infection in goats, each dominant VSG was represented by several closely related transcripts displaying comparable variation to what we now see within gene arrays (Silva-Pereira et al 2020). It is unclear if this was reproduced here. The P151 array contains 27 intact VSG genes with between 0.003-0.053% nucleotide divergence, translating into an average of 19 amino acid substitutions in pairwise comparison. When transcript abundance is measured with FeatureCounts on ONT RNA reads (Table S2), TvY486_LIV26_0061645 is the most abundant transcript (count = 37850), but all tandem copies are expressed, and six non-identical genes have an abundance of similar magnitude (counts between 13802 and 35297). However, this could reflect mappings of reads from one transcript to multiple, near-identical loci; when RNA-seq reads are mapped to P151 gene sequences and abundance measured with Kallisto, only TvY486_LIV26_0061645 is abundant (TPM= 2699.6); 25 other P151 genes are measured but none have a TPM >1.5. We assembled P151 transcripts de novo from Novaseq data using Trinity and measured their abundance with Kallisto (Table S2). This produced four distinct sequences, the most abundant (DN22938_021) was again derived from TvY486_LIV26_0061645. Finally, we assembled ONT RNA reads with RATTLE v1.0 (de la Rubia et al. 2022); all P151 sequences in this assembly were identical to TvY486_LIV26_0061645. Therefore, it is unclear if multiple transcripts are expressed from the active VSG gene array or not, but it seems likely that, the tandem gene arrangement notwithstanding, one paralog contributes most of the active VSG transcripts.

For comparison, the analysis was repeated using RNA-seq data from two previous *T. vivax* blood infections. Fig. 2e shows that a superabundant VSG transcript is again evident in a mouse infection with the same Y486 strain (carried out by Jackson et al. (2015)), but it is P121, not P151 (neither P151 nor P36 were transcribed in the 2015 experiment). Another analysis of VSG expression in goats using a different *T. vivax* strain (Lins) also displays a superabundant VSG (P174; Fig. 2g), but it is different to both Y486 mouse experiments.

It is worth highlighting the expression of vivaxin in Fig. 2, a *T. vivax*-specific gene family that is predicted to be expressed alongside VSG on the bloodstream-stage cell surface. TvY486_LIV26_0029790 is a beta-vivaxin gene ranked first in transcript abundance (by FeatureCounts) and 11^th^ (by Kallisto) across all genes. Its transcript (DN1908_911; TPM=2355.7) is approximately equal to P151 (DN22938_021; TPM=2160.7) in abundance in this study. This is same sequence as the highly abundant ‘antigen-4’ (vivaxin-β8) observed in goats infected with *T. vivax* Lins (Romero-Ramirez et al. 2022). Depending on the measure chosen, other vivaxin genes are also strongly transcribed also; eight of the 25 most abundant transcripts derive from vivaxin genes when ranked by FeatureCounts (Table S2). Taking all Novaseq RNA-seq reads together, 3.1% map to vivaxin and a further 3.1% map to other *T. vivax*-specific cell surface genes (Fig. 2b), proportions that remain roughly consistent on previous occasions (Fig. 2d,f).

Proteomics analysis, carried out to further probe VSG expression, confirmed the dominance of P151 and P36. LC-MS identified 805 proteins, including 21 VSG derived from 10 gene arrays, as described in Table S3. Table 1 describes 15 VSG genes for which >1 peptides were observed. As with the transcriptomic analysis, TvY486_LIV26_0061645 (P151) was the most abundant VSG protein by peptide count, while P36 was also expressed. However, five other VSG proteins (P34, P89, P121, P147, P161) were observed although none displayed abundant transcripts, despite the RNA and protein samples being prepared from the same cell isolations. Note that P121, the superabundant VSG in the 2015 transcriptome, was expressed as a minor VSG here, suggesting that some cells retained that epigenetic condition. In contrast to the ambiguity in the transcriptome, Table 1 shows that peptides mapped uniquely to three different P151 genes.

VSG expression is summarised in Fig. 3. The active VSG, P151, is encoded from a single, sub-telomeric gene array (Fig. 3a). We cannot say presently that all arrays are available for activation; there may be structural requirements for activation that are not apparent from our data. Some other minor variants are present among both transcripts and peptides, which may represent rare clones in the population (e.g. P121), but, nonetheless, Fig. 3b illustrates how the VSG array adjacent to P151 was transcriptionally silent, as were almost all other arrays. Noting that previous studies also displayed dominant, superabundant VSG transcripts but that these belonged to different phylotypes, this indicates that *T. vivax* displays monoallelic expression and antigenic switching at the genomic level. Fig. 3c shows how multiple peptides mapped to the active VSG, providing 44% coverage of the coding sequence. Note also the expression of the *ingi* element upstream of the P151 array in Fig. 3b. Perhaps an active VSG array is expressed as a polycistronic transcript from a promoter in the 5’ region, and thus VSG expression in *T. vivax* might utilise the same regulatory mechanism as other genes.

**Fig. 3.**
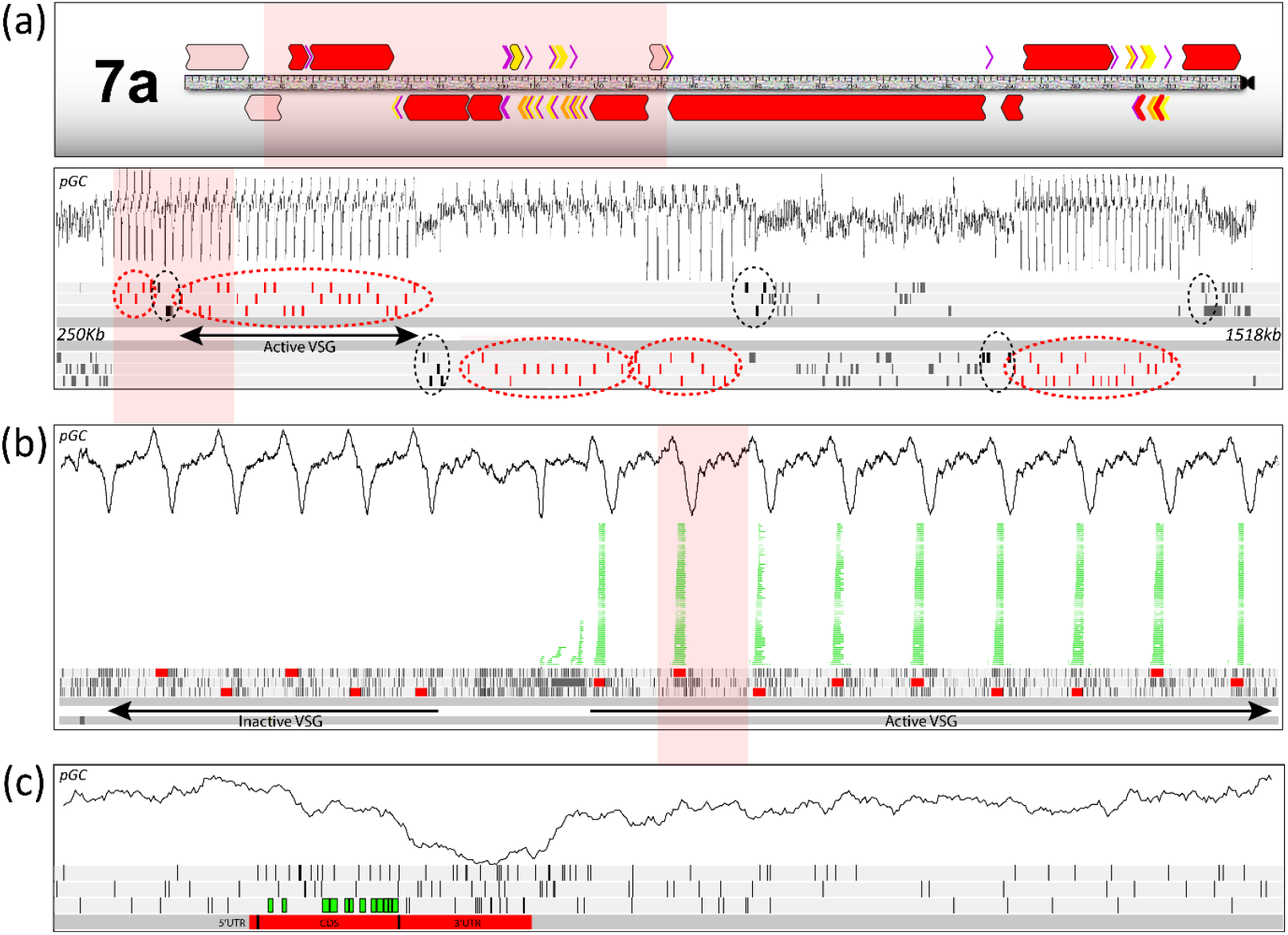
VSG expression from sub-telomeric gene arrays. Expressed VSG were detected in a mouse bloodstream infection using proteomic analysis. The genomic context of the most abundant (‘active’) VSG protein (P151, array 88, e.g. TvY486_LIV26_0061645) is shown. **(a)** Part of scaffold 7a (250-1,518Kb inclusive) showing proportion of GC (*pGC*), forward and reverse DNA strands (dark grey) and six coding frames (light grey). Rectangles with reading frames denote VSG (red) or other coding sequences (grey). Dotted lines denote the position of VSG gene arrays (red) or *ingi* transposable elements (black). **(b)** ONT RNA reads (green) are mapped by blastn to the portion of (a) indicated in pink; multiple mapping was allowed. **(c)** Peptides observed by mass spectrometry (green boxes) are mapped to the portion of (b) indicated in pink; unique peptides were detected from at least three gene copies.

Hence, while *T. vivax* shares an antigenic variation phenotype with other African trypanosomes, its genome shows that the regulatory mechanism for monoallelic expression and antigenic switching must be different from the model established for *T. brucei*. That model may reflect a level of sophistication in *T. brucei* that is highly derived and absent in other species.

## Conclusions

The 69.86 Mbp *T. vivax* Y486 assembly consists of 23 scaffolds (69,756,071 bp), a mitochondrial genome (32,130 bp), and 53 unassembled bin contigs (108,174 bp). Like other African trypanosomes, the chromosomes combine polycistronic cores that likely reflect fusion events in the evolutionary past with hemizygous sub-telomeres that contain VSG gene arrays, separated by transposable elements. In these aspects, *T. vivax* conforms to the African trypanosome norm, but the detail shows fundamental differences with other species. First, *T. vivax*-specific cell surface genes are concentrated in two sub-telomeres; these include at least one non-VSG surface protein (β-vivaxin) that is comparable in abundance to VSG in bloodstream-stage cells. Second, the VSG are mostly intact genes, arranged in tandem arrays of near-perfect repetition, each array being the unique location of just one antigen. Third, transcriptomic and proteomic evidence shows that VSG are expressed from sub-telomeric polycistrons, and there is no evidence for telomeric expression or any dedicated expression site. In these and other respects, *T. vivax* departs from the physiological models established for *T. brucei*; we have presented an improved genome assembly that provides a robust basis to exploring this important pathogen’s unique biology.

## Methods

### Ethics statement

All procedures involving animals were carried out under the authority of a UK Home Office Project Licence in accordance with the ‘Animals (Scientific Procedures) Act 1986’ and Directive 2010/63/EU and were approved by The University of Edinburgh Bioresearch & Veterinary Services Animal Welfare and Ethics Review Body (AWERB). Animals were maintained in a standard 12-h light-12-h dark cycles and given access to food and water *ad libitum*.

### DNA sequencing

*Trypanosoma vivax* Y486 epimastigote-stage cells were grown *in vitro* as described previously (D’Archivio et al. 2011) and high molecular weight genomic DNA was prepared from approximately 1x10^9^ cells using the Circulomics Nanobind CBB kit using the UHMW protocol. The same DNA preparation was sequenced on three platforms. First, HiFi reads were produced using the PacBio Sequel II SMRT cell in CCS run mode. Second, paired-end 150 bp reads were generated using the Illumina NovaSeq X Plus with 10B chemistry. Third, nanopore (‘ONT’) reads were produced on the Oxford Nanopore GridION. In addition, DNA was extracted from an independent cell preparation of the same Y486 strain for high-throughput chromosome conformation capture (Hi-C). The Hi-C library was generated from approximately 7 × 10^8^ cells using the Proximo Hi-C (Microbe) Kit as per manufacturer’s instructions.

### Genome assembly

The genome sequence was assembled from HiFi reads scaffolded with Hi-C and polished using Illumina data. HiFi reads were first filtered for residual adapter sequence using hiFiAdpaterFilt v1 (Sim et al. 2022) and then assembled using hifiasm v0.19 (Cheng et al. 2021). Quality was assessed with quast v5.0.2 (Gurevich et al. 2013) and BUSCO v5.2.2 (Manni et al. 2021). Potential contamination was examined using blobtools v2 (Laetsch & Blaxter 2017) and none was found. Two rounds of haplotig purging were performed with purge_dups v1.2.5 (Guan et al. 2020) to remove any heterozygous duplication present in the assembly. The purged assembly was scaffolded using Hi-C with YaHS v1.2a.2 (Zhou et al. 2023); JBAT v1.11.08 (Durand et al. 2016) was used to visualise Hi-C contacts. A second assembly was attempted by combining HiFi and ONT reads using hifiasm v0.19, followed by two rounds of haplotig purging and then scaffolding with Hi-C data as before. The assembled scaffolds were compared to one another using BLASTn, which showed that there were perfect sequence overlaps between nine pairs of scaffolds; these overlapping ends were articulated. Illumina data were assembled separately using SPAdes (Prjibelski et al. 2020) to check for very small DNA species that may have been omitted due to size-selection by long-read sequencing and assembly.

### Genome annotation

The existing Y486 genome annotation was ported to the HiFi-Hi-C assembly with COMPANION v2.2.8 (Haese-Hill, Crouch & Otto 2024). Since this was likely to be affected by the abundant sequence gaps in the original assembly, all protein-coding gene models were then manually annotated. Sequence homology with *T. brucei* and *T. cruzi* coding sequences was inferred using tBLASTn and significant matches (i.e. e-value > 10e^-4^) were mapped to the new assembly. RNA-seq data from all three *T. vivax* life stages that were published previously (Jackson et al. 2015) were mapped to the new assembly using Hisat2 (Kim et al. 2019). This evidence, combined with assembled transcripts from RNAseq in this study (see below), was used to confirm or reject putative coding sequences, refine gene model boundaries and apply product descriptions. To annotate homoeology between *T. vivax* and *T. brucei* chromosomes, the new assembly was aligned with the 11 chromosome *T. brucei* assembly (TritrypDB v69) using ProgressiveMAUVE (Darling, Mau & Perna 2010).

### VSG repertoire

VSG sequences were identified using tblastn; the query in these searches were 178 amino acid sequences that represent all VSG ‘phylotypes’ that we previously described (Silva-Pereira et al. 2020). tBLASTn matches with a minimum of 90% sequence identity and 1,500 bp in length were selected and mapped to the genome annotation. The highest match for a given VSG gene was used to ascribe a phylotype to each. It became clear that the new assembly contained VSG genes that had not been examined previously and these were made into additional phylotypes.

### Measurement of VSG expression

Ten-week-old CD1 mice were injected subcutaneously with approximately 10^5^ in vitro-derived metacyclic-like cells from the same *T. vivax* Y486 stabilate that had been used in DNA preparation. Briefly, metacyclic-like *T. vivax* parasites were enriched from epimastigote cultures, using complement-mediated lysis of epimastigotes. Parasites were incubated for 4 hours at 37 °C in medium containing 50% non-heat inactivated foetal bovine serum, with occasional mixing. Epimastigote ‘clumps’ were removed from the cell suspension using a 5 µm filter. The resulting enriched suspension was pelleted at 1800 × *g*, then resuspended in TV3 medium for injection into mice. Mice were weighed, condition-scored and bled daily to determine parasitaemia counts as described previously (Herbert & Lumsden 1976). Once parasitaemia reached 5x10^8^ parasites/ml, blood was collected by cardiac puncture in heparin (2500 IU/kg) and processed as described previously (D’Archivio et al. 2011). RNA was isolated from the resulting *ex vivo* bloodstream-form *T. vivax* parasites from two biological replicates by Trizol extraction (Thermo Fisher Scientific), followed by RNeasy Plus Mini Kit clean-up (Qiagen), as per manufacturer’s instructions. Cell lysate from another biological replicate was prepared for proteomics in RIPA lysis buffer (Life Technologies) plus HALT protease inhibitor cocktail (VWR).

For transcriptomic analysis, each parasite RNA sample was sequenced on the Illumina Novaseq platform and the RNA-seq reads were mapped to the new assembly using Hisat2. Transcript abundance was estimated for all annotated coding sequences using Kallisto (Bray et al. 2016) and FeatureCounts (Liao, Smyth & Shi 2014) and averaged over the two replicates. To check for unannotated features, transcript sequences were assembled using Trinity (Grabherr et al. 2011) and validated by mapping back to the genome sequence with blastn. For those transcripts mapping to the genome (≥99% sequence identity), sequence abundance was estimated by mapping RNA-seq reads back to the transcriptome assembly with Kallisto. For comparison, data from two previous *T. vivax* bloodstream infections in mouse (Jackson et al. 2015) and goat (Silva-Pereira et al. 2020) were also mapped to the new assembly and quantified.

For proteomic analysis, parasite protein was analysed by Liquid Chromatography-Mass Spectroscopy (LC-MS). After denaturation and centrifugation at 600 RPM (40 × *g*), the sample was quantified using nanodrop with an average reading of 21.2 mg/mL. Following cysteine reduction by adding 4 mM DTT solution (60°C for 10 min with 40 × *g*) and alkylation with 7 mM iodoacetamide (30 min), 120 ng of SP3 beads were added, followed by a total volume of 80% Acetonitrile (30 min). After washing, the beads were dried and resuspended in 50 mM AmBic. In-solution digestion was carried out by adding 10 ng of trypsin and incubating at 37°C for 16 h with 1500 rpm (250 × *g*). 6 μL of digested protein was analysed using an Ultimate 3000 RSLC™ nano-system (Thermo Scientific, Hemel Hempstead) coupled to an Orbitrap Exploris 480™ mass spectrometer (Thermo Scientific). Analysis of the LC-MS/MS data was performed using FragPipe V 23.1 (Hsiao et al. 2024). Data were searched using Mascot against the UniProt *Trypanosoma vivax* (strain Y486) database, (download 16-1-2026). The FragPipe contaminant database was added to the search to identify known contaminants.

## Supplementary Material

**Fig. S1**. Hi-C contact map for the Hifi-Hi-C hybrid assembly. Topological associated domains coincide with 29 contigs in the original assembly. Further manual articulations to combine contigs with sequence overlap produced 24 scaffolds (as shown in Figure 1). The correspondence between contigs and scaffolds is shown at right.

**Fig. S2**. Hi-C contact map for the ONT-Hi-C hybrid assembly. Topological associated domains coincide with 29 contigs in the ONT-Hi-C assembly. Three contigs contained internal stop codons (black arrows). The correspondence between ONT-Hi-C contigs and Hifi-Hi-C contigs is shown at right. Note that several ONT-Hi-C contigs (e.g. #12) combine several separate contigs from the Hifi-Hi-C assembly that have telomeres attached (i.e. #1,5,6,9).

**Fig. S3**. A dispersed segmental duplication that is not associated with VSG genes. The figure depicts the largest (and only substantial) duplicon in the *T. vivax* Y486 genome, discovered by mapping all contigs to one another. The duplicon concerns a 40 kb region of scaffold 7 that includes around 20 genes (pink boxes); these are part of a conserved polycistron on chromosome 11 in *T. brucei*. The duplicon is copied at 16 other locations (grey bars), with variable size and completeness, and often positioned between VSG gene arrays. Seven of the duplicated genes are be linked to ciliary/flagella function by gene ontology: octanoyltransferase (TvY486_LIV26_0056160); TFIIH basal transcription factor subunit (TvY486_LIV26_0056180); Component of motile flagella 63 (TvY486_LIV26_0056185); cyclophilin (TvY486_LIV26_0056190); ADP-ribosylation factor GTPase activating protein (TvY486_LIV26_0056205); tubulin binding cofactor c (TvY486_LIV26_0056210); CobW/HypB/UreG, nucleotide-binding domain containing protein (TvY486_LIV26_0056215).

**Table S1**. VSG gene arrays in the *T. vivax* Y486 LIV26 genome assembly.

**Table S2**. RNA-seq for a *T. vivax* Y486 LIV26 blood infection in a mouse model, showing transcript abundance calculated with both FeatureCounts and Kallisto.

**Table S3**. Peptides detected by mass spectrometry analysis of whole protein prepared by a *T. vivax* Y486 blood infection in mouse.

**Table S4**. Accession numbers for sequence read data deposited in the European Nucleotide Archive.

## Acknowledgements

We thank Philip Brownridge of the University of Liverpool Centre for Proteomic Research for assistance with proteomic analysis. Hi-C library sequencing was performed by the University of Edinburgh Genetics Core, Edinburgh Clinical Research Facility (RRID:SCR_025433). We thank Heather Warnock, Scott McVittie and Lee McManus, from the Biological Research Facility at The Roslin Institute, University of Edinburgh for their excellent technical support. This work was supported by the Wellcome Trust (206815/Z/17/Z to A.P.J. as co-investigator), and the BBSRC (BB/X009807/1 to P.C.S. and BB/W000296/1 to P.C.S. and L.J.M.). The Roslin Institute is supported by core funding from the BBSRC (BBS/E/RL/230002C).

## Author Contributions

A.P.J., R.Y., M.A.W., P.C.S. and L.J.M. wrote and edited the manuscript. A.P.J., R.Y., M.A.W. and P.C.S. were responsible for methodology, investigation and analysis. A.P.J., A.C.D. and L.J.M. supervised and conceptualized the project, and secured funding.

## Data Availability

The final genome assembly is deposited in Genbank with accession GCA_982375215 (Bioproject PRJEB108666). All raw sequence data relating to genome and transcriptome sequencing is deposited in the European Nucleotide Archive; accession numbers are provided in Table S4. The mass spectrometry proteomics data have been deposited to the ProteomeXchange Consortium via the PRIDE partner repository with the dataset identifier PXD075690 and 10.6019/PXD075690.

