## Supplementary Material for "An improved *Trypanosoma vivax* genome assembly confirms the distinct organisation of its variant antigen repertoire"

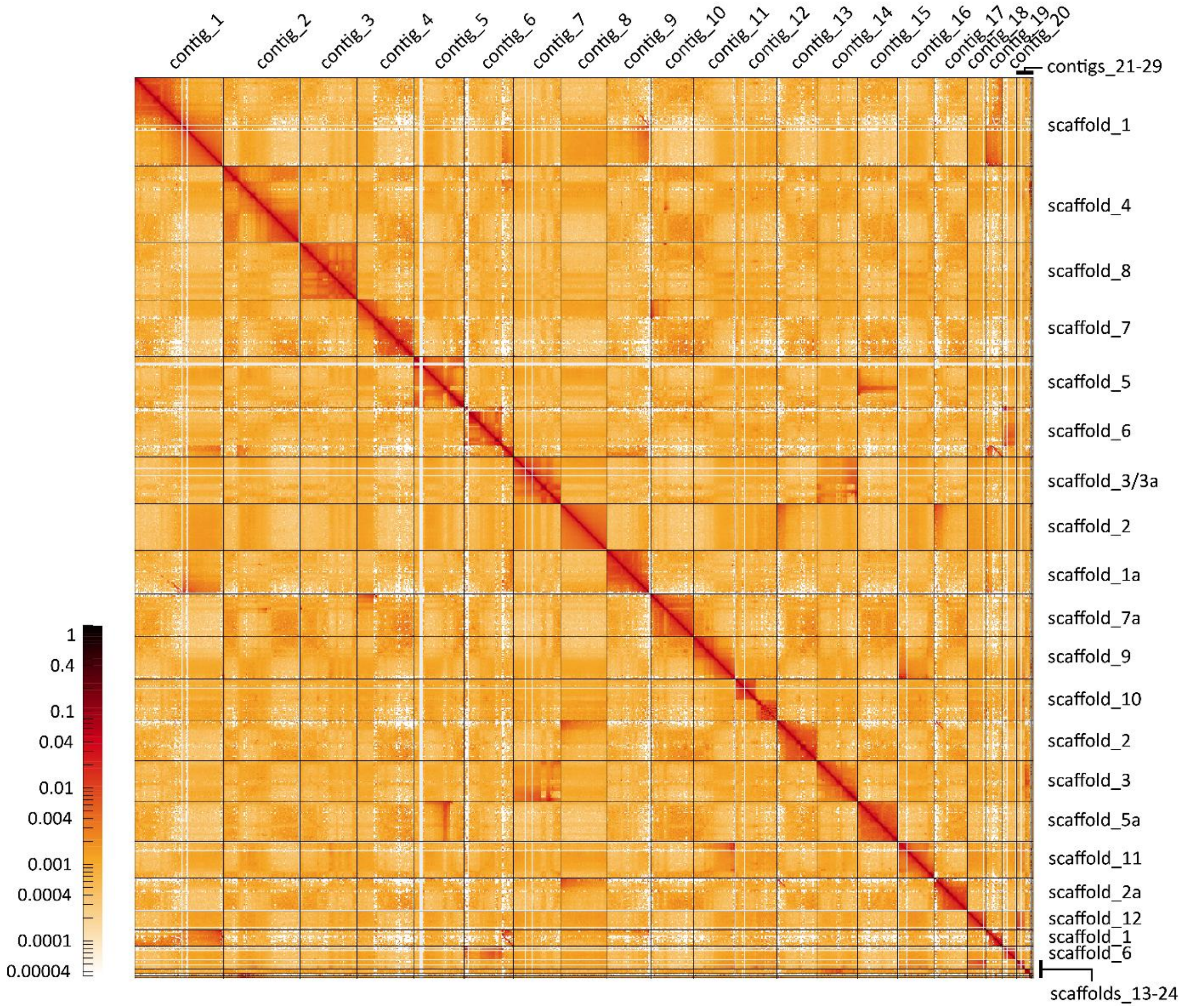


**Fig. S1.** Hi-C contact map for the Hifi-HiC hybrid assembly. Topological associated domains coincide with 29 contigs in the original assembly. Further manual articulations to combine contigs with sequence overlap produced 24 scaffolds (as shown in Figure 1). The correspondence between contigs and scaffolds is shown at right.

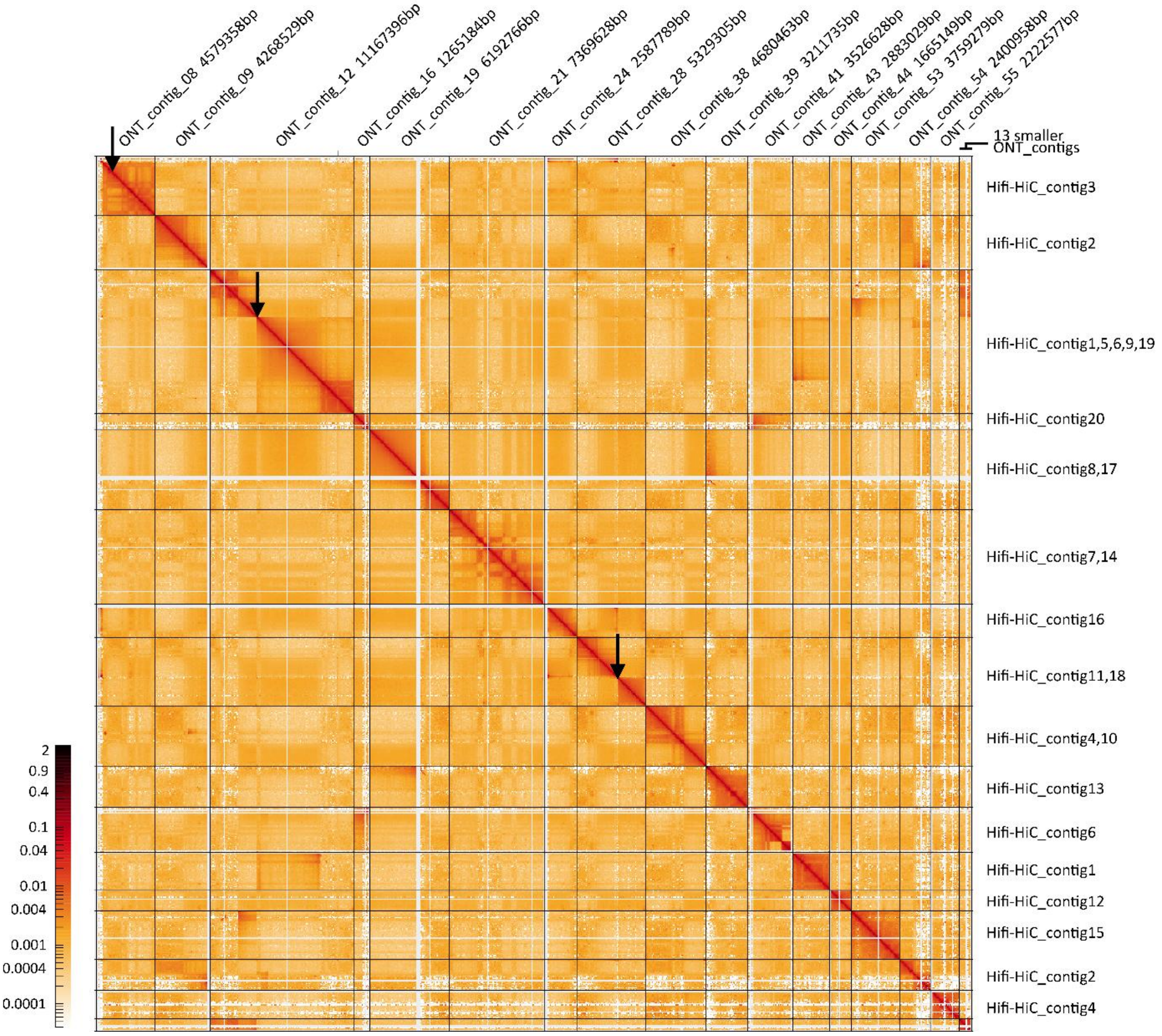


**Fig. S2.** Hi-C contact map for the ONT-HiC hybrid assembly. Topological associated domains coincide with 29 contigs in the ONT-HiC assembly. Three contigs contained internal stop codons (black arrows). The correspondence between ONT-HiC contigs and Hifi-HiC contigs is shown at right. Note that several ONT-HiC contigs (e.g. #12) combine several separate contigs from the Hifi-HiC assembly that have telomeres attached (i.e. #1,5,6,9).

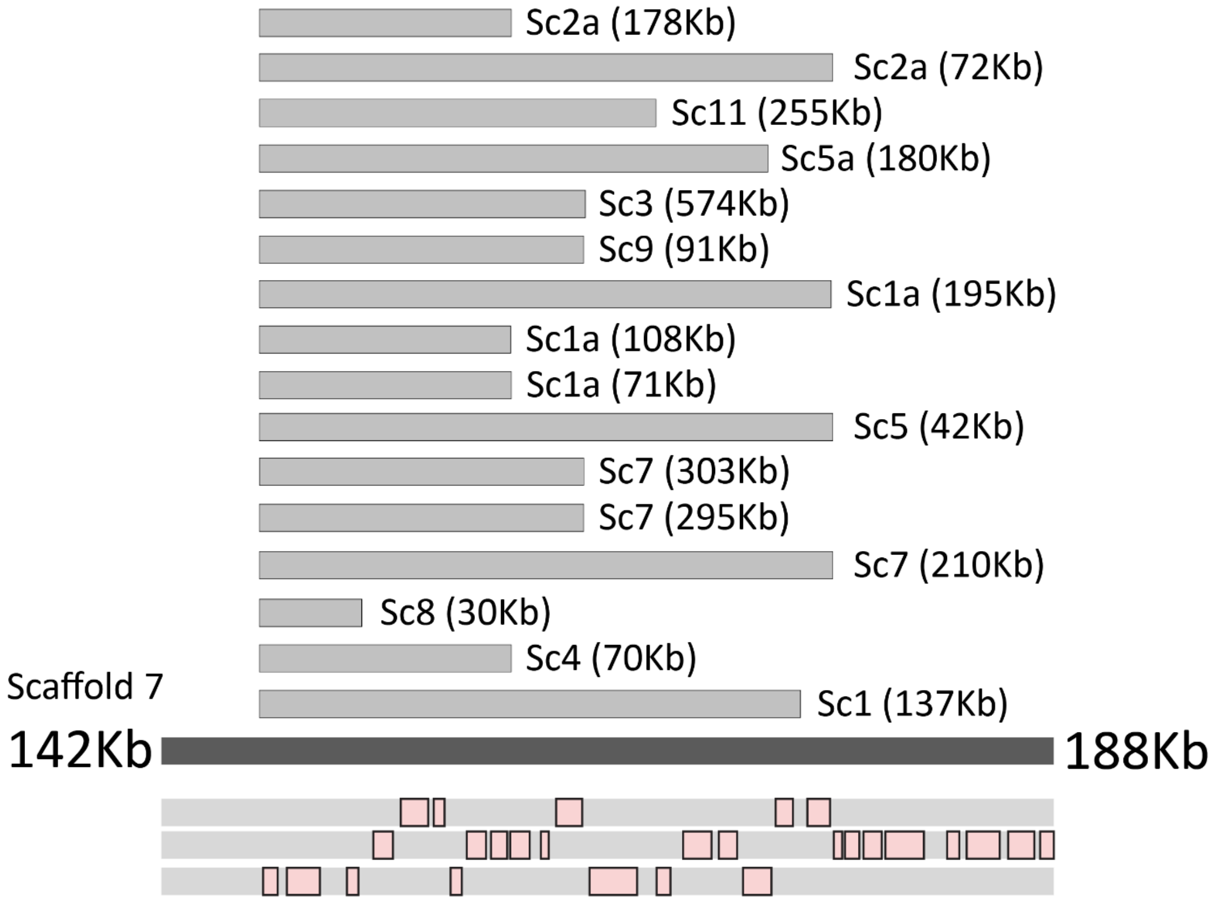


**Fig. S3.** A dispersed segmental duplication that is not associated with VSG genes. The figure depicts the largest (and only substantial) duplicon in the *T. vivax* Y486 genome, discovered by mapping all contigs to one another. The duplicon concerns a 40kb region of scaffold 7 that includes around 20 genes (pink boxes); these are part of a conserved polycistron on chromosome 11 in *T. brucei*. The duplicon is copied at 16 other locations (grey bars), with variable size and completeness, and often positioned between VSG gene arrays. Seven of the duplicated genes are be linked to ciliary/flagella function by gene ontology: octanoyltransferase (TvY486_LIV26_0056160); TFIIH basal transcription factor subunit (TvY486_LIV26_0056180); Component of motile flagella 63 (TvY486_LIV26_0056185); cyclophilin (TvY486_LIV26_0056190); ADP-ribosylation factor GTPase activating protein (TvY486_LIV26_0056205); tubulin binding cofactor c (TvY486_LIV26_0056210); CobW/HypB/UreG, nucleotide-binding domain containing protein (TvY486_LIV26_0056215).
